# Picoeukaryotic photosynthetic potential is functionally redundant but taxonomically structured at global scale

**DOI:** 10.1101/2023.09.22.558943

**Authors:** Alexandre Schickele, Pavla Debeljak, Sakina-Dorothée Ayata, Lucie Bittner, Eric Pelletier, Lionel Guidi, Jean-Olivier Irisson

**Author notes:** These authors contributed equally (Lionel Guidi, Jean-Olivier Irisson).

## Abstract

Primary production, performed by RUBISCO, and often associated with carbon concentration mechanisms, is of major importance in the oceans. Thanks to growing metagenomic resources (e.g., eukaryotic Metagenome-Assembled-Genomes; MAGs), we provide the first reproducible machine-learning-based framework to derive the potential biogeography of a given function, through the multi-output regression of the standardized number of reads of the associated genes on environmental climatologies. We use it to study the genomic potential of C4-photosynthesis of picoeukaryotes, a diverse and abundant group of marine unicellular photosynthetic organisms. We show that the genomic potential supporting C4-enzymes and RUBISCO exhibit strong functional redundancy and an important affinity towards tropical oligotrophic waters. This redundancy is then structured taxonomically by the dominance of Mamiellophyceae and Prymnesiophyceae in mid and high latitudes. Finally, unlike the genomic potential related to most C4-enzymes, the one of RUBISCO showed a clear pattern affinity for temperate waters.

## INTRODUCTION

Most of the photosynthetic production on earth relies on the ribulose-1,5-bisphosphate carboxylase oxygenase (RUBISCO; 1). However, because RUBISCO emerged *∼*2 billion years ago in a period characterized by low oxygen (2), its carboxylase function is surprisingly inefficient relative to its oxygenase function, when considering the contemporary CO_2_-to-oxygen ratio (3). To compensate for this metabolic caveat related to RUBISCO-only photosynthesis (i.e., C3-photosynthesis), carbon fixation pathways evolved *∼*30 million years ago, when atmospheric CO_2_ levels were estimated under 200 ppm. The latter induced selective pressure towards higher carbon fixation efficiency, leading to the development of various Carbon Concentration Mechanisms (CCMs; i.e., biophysical or biochemical) to compensate for the photorespiration affinity of RUBISCO (4). Among biochemical CCMs, C4-enzymes independently evolved across a large variety of marine and terrestrial lineages (4, 5). The C4 cycle is performed through 3 acid-decarboxylation types, leading to an increase of the CO_2_-to-oxygen ratio at the active site of RUBISCO (6): the MDC-NADP type, the MDC-NAD type, and the PEPCK type. The common enzyme to all C4 acid decarboxylation types is phosphoenolpyruvate carboxylase (PEPC), fixing CO_2_ in the cytosol by producing oxaloacetate. In the MDC-NADP type, oxaloacetate is transferred to the chloroplast and reduced to malate. The latter is then decarboxylated, producing CO_2_ and pyruvate, which is converted back to phosphoenolpyruvate. In the MDC-NAD type, oxaloacetate is transferred to the mitochondria and reduced to malate. The decarboxylation reaction transfers CO_2_ to the chloroplast by producing pyruvate that is transferred back to the chloroplast to be converted to phosphoenolpyruvate. Finally, the PEPCK type directly converts the mitochondrial oxaloacetate to phosphoenolpyruvate. However, it partially performs the MDH-NAD reduction and MDC-NADP decarboxylation reactions to balance the ATP and NADPH budget, leading to common reactions and enzymes between acid-decarboxylation types (6). In the terrestrial realm, both physiological measurements and stable isotope techniques confirmed the presence of C3-photoynthesis across a large range of environmental conditions, conversely to C4-photosynthesis that is adapted to warm, nutrient poor and high irradiance conditions (7, 8). In the marine realm however, only a few studies explored the environmental affinity of C4-photosynthesis regarding terrestrial-based hypotheses (e.g., 5, 9, 10). The potential for C4-photosynthesis is highly suspected in key picoeukaryote lineages such as Mamiellophyceae and Prymnesiophyceae. Currently, subcellular evidence for C4-enzymes include (i) MDC-NADP and PEPC in *Ostreococcus Tauri* (11), (ii) MDC-NADP, PEPC, three different oxoglutarate-to-malate translocator and pyruvate phosphate dikinase (PEPDK) in various Micromonas strains (12) and (iii) PEPC in Prymnesiophyceae (Emiliana Huxleyi; plastid presence and gene encoding;, 13).

Marine carbon fixation is largely performed by picoeukaryotes (e.g., 30 to 50 % of global primary production, 14, 15), some of which are suspected to use C4-photosynthesis (e.g., in picoeukaryotic diatoms;, 5, 9). Picoeukaryotes correspond to the unicellular eukaryotic marine plankton, that are among the most diverse and abundant organisms in the sunlit layer of the world ocean (16–18). In nutrient-poor areas, such as the oligotrophic open ocean, they locally contribute up to 80 % of the phytoplanktonic biomass (19). However, because of their size (i.e., 0.8 to 5 μm), poor representation in culture collections (20) and thus the difficulty for both physiological measurements and stable isotope analysis in natural populations, the genomic potential supporting C3, and C4-photosynthesis, its associated biogeography and functioning remains scarcely documented (5, 8, 9).

Recent global expeditions focusing on surface plankton sampling, together with advances in metagenomic sequencing, provided unique data to address the genomic potential and biogeography-related gaps (e.g., 21–24). In this context, metagenomics data are of growing interest to explore the hidden taxonomic and functional diversity potentially related to carbon fixation in picoeukaryotes (e.g., 25, 26). For example, genome-resolved metagenomics (27) based on the *Tara-*Oceans eukaryotic metagenome led to the reconstruction of ∼800 Metagenome-Assembled-Genomes (MAGs; 28). The latter are defined as genome-based taxonomic units, functionally and taxonomically annotated, and quantified by their associated genome-wide metagenomic reads. Therefore, MAGs offer the unique opportunity to study the genomic potential supporting carbon fixation and its biogeography, through both a functional and a taxonomic prism.

Habitat modelling is a popular niche theory-based tool to estimate species biogeography according to the environmental conditions in which they are observed (29). Marine organisms are known for their important sensitivity to their surrounding environmental conditions, influencing growth, reproduction, and metabolic efficiency across all life stages (30). Thus, habitat modelling has been widely used to project the past, present, and future biogeography across various marine organisms, from zooplankton to fishes (e.g., 31). However, omics-based habitat modelling is still an emerging field to explore functional and taxonomic biogeography associated with unicellular planktonic organisms (32–34). Building on the above-mentioned properties associated with MAGs, habitat modelling is transferable to genomic potential, thus exploring the quantitative response of the associated taxonomic and functional gene annotations to environmental conditions.

Here, complementing recent studies on prokaryote - environment relationships (32), we provide an original, machine learning-based, comprehensive, and reproducible framework to derive the biogeography of the genomic potential related to metabolic functions, from metagenomic-based relative abundances data. Using Multivariate Boosted Tree Regressors (35), we simultaneously project the biogeography of selected genomic functional annotations, while accounting both for their interactions and environmental responses. We applied this framework to metagenome-based Protein Functional Clusters (PFCs; hereafter referred to as “clusters”) linked to RUBISCO and C4-enzymes only, in marine picoeukaryotes. Compared to a more traditional approach (i.e., searching reads in a functional database using sequence similarity), our methodology combining MAGs and PFCs offers several advantages. The quantitative signal resulting from a MAG is (i) standardized by the genome length and (ii) correspond to a taxonomic identity. Combined in PFCs, (iii) it also includes the fraction of signal corresponding to not yet annotated genes. Thus, this approach offers a more robust quantitative framework than traditional approaches, representative of eukaryotic plankton diversity in open oceans (39.1 billion reads recruited, ∼97% identity, ∼25 Gbp;, 28) and transferable to a variety of functions or enzymes of interest using the already computed PFC network. Finally, habitat modeling provides an interesting tool to estimate the response and co-dominance patterns of C4-enzymes and RUBISCO to environmental conditions representative of the global ocean, conversely to estimates from the samples only, that might be driven by sampling and associated environmental biases.

## RESULTS

### 2.1. C4-CCM enzymes across sampled stations

From the *Tara Oceans* eukaryotic MAGs, ∼1.2 million clusters were built, for which 349 are related to RUBISCO or C4-enzymes (**Fig. S1, Table S1**). This dataset corresponds to 817 unique genes, with a median observed presence across 45 sampled stations per cluster. To avoid considering enzymes related to other metabolic functions, we only selected those related to RUBISCO or C4-enzymes only, corresponding to 240 clusters, distributed across the world Ocean except the Arctic, western Pacific and to a lesser extent Southern Ocean (**Figure 1**). The successive cluster selection criteria (i.e., PFCs exclusive to RUBISCO or C4-enzymes, minimum presence at 10 sampling stations) did not present significant effects on the distribution of clusters across number of reads, number of genes and taxonomic classes (**Fig. S3)**. In contrast, we observed a loss of signal for the MDCs (-NAD and -NADP), between functionally exclusive and non-exclusive clusters, highlighting an important fraction of sequence homologs for these enzymes (**Fig. S3**).

**Figure 1.**
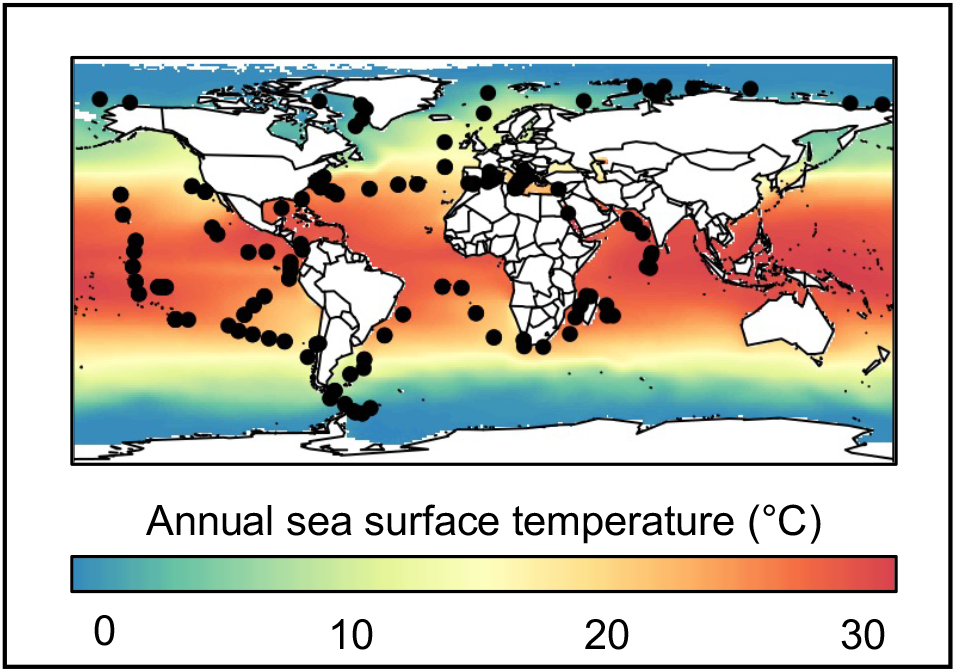
Location of the *Tara Oceans* (TO) sampling stations, represented as black dots. Annual sea surface temperature from World Ocean Atlas (Boyer et al. 2018) are represented in background.

### 2.2. Standardized distribution of the genomic potential related to C4-photosynthesis

Here we present projections for each C4-enzyme and RUBISCO. First, we rescaled the cluster-level projections (i.e., model outputs; **Fig. S1D**) between 0 and 1 (i.e., distribution patterns, **Fig. S2**). Then, we aggregated these patterns at the enzyme-level according to their respective functional annotation. We therefore alleviated the propagation of the observed dominance of a given cluster to the aggregated enzyme-level patterns. The resulting enzyme-level projections are referred to as standardized patterns. For each enzyme, it represents a prediction of the genomic potential according to the environmental conditions at each geographical location, and independently of any taxonomic dominance.

Because most C4-enzymes are involved in several acid-decarboxylation types, we cannot directly infer their corresponding distribution. However, MDC - NAD, MDC - NADP and PEPCK are considered representative of their respective acid-decarboxylation types. We predicted similar standardized patterns (**Figure 2)** for all acid decarboxylation types and RUBISCO. The standardized patterns of all C4-enzymes presented medium to high pairwise Pearson’s correlation (0.5 to 0.9), except MDC - NAD and GOT which are weakly correlated (0.3).

**Figure 2.**
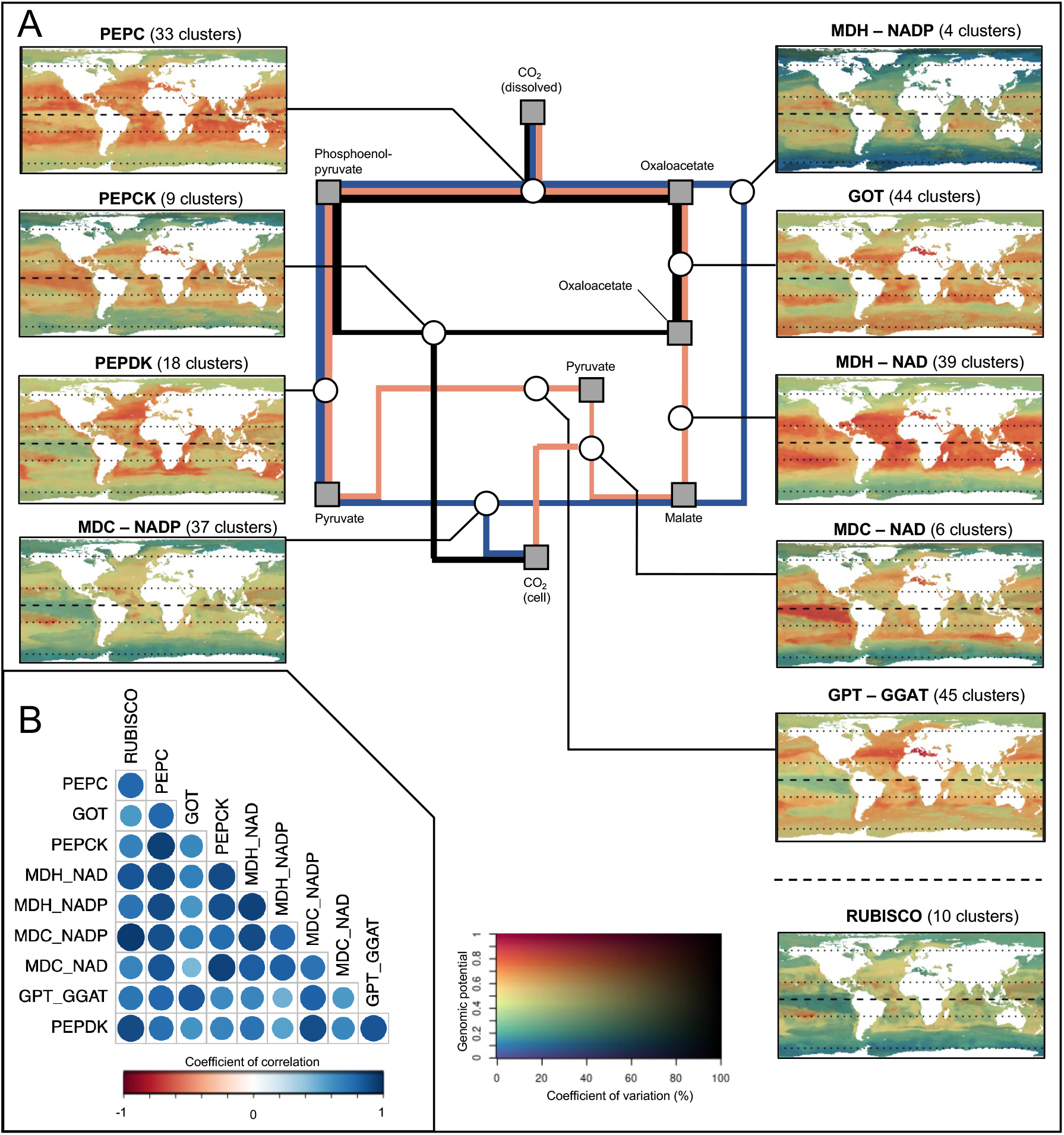
Standardized patterns corresponding to the relative genomic potential supporting C4-enzymes and RUBISCO. **(A)** Synthetic diagram of the metabolic pathway and corresponding projections. **(B)** Inter-projections Pearson’s spatial correlation index. The three mains currently described acid-decarboxylation types are represented in blue (MDC-NADP), red (MDC-NAD) and black (PEPCK), respectively. Involved metabolic components and enzymes are indicated on the diagram by squares and circles, respectively. The 2D color scale represents the standardized genomic potential for the target enzyme as the hue value (Y-axis) and the associated coefficient of variation as the saturation (i.e., uncertainty in % of the mean; X-axis). An orange to red hue corresponds to region where environmental conditions yield a high proportion (>0.6) of the target genes in the model. A low saturation level corresponds to an important variance among the underlying cluster-level projections.

We predicted a high genomic potential (> 0.6) for all standardized patterns in temperate to tropical latitudes, with an associated coefficient of variation below 30 % (**Figure 2A**). We also predicted a high potential (> 0.8) for RUBISCO and PEPDK for temperate to tropical waters only. In contrast, the potential for PEPC, GOT, MDCs and MDHs were high in equatorial latitudes. These patterns suggest a higher affinity of the genomic potential of C4-enzymes for the equatorial ocean, in comparison to RUBISCO. Furthermore, we predicted low-to-moderate potential (between 0 and 0.4) in high latitudes (i.e., above polar circles) for all standardized patterns (**Figure 2A**). Predictions in such latitudes also present important calibration and projection-related variability, with coefficients of variations ranging from 30 to 100 % (e.g., for the MDH – NADP and PEPCK). Therefore, our genomic potential predictions remain inconclusive in high latitudes, also subject to lower sampling coverage.

The environmental variables importance in the trained model (**Fig. S4**) highlighted the predominant roles of dissolved oxygen concentration (contributing to 34% of the explained variance) and the yearly variability (i.e., inter-month standard deviation) in Salinity (29%) and, to a lesser extent, of oxygen saturation, chlorophyll a concentration and temperature. Furthermore, we revealed a strong affinity (i.e., maximum potential) of most standardized patterns (**Fig. S5**) for tropical, oligotrophic conditions (e.g., temperature between 15 to 30 °C; phosphate concentration below 0.5 μmol/kg). However, we predicted different responses to the variability in Chlorophyll a concentration and euphotic zone depth across enzymes (**Fig. S5**). Finally, we highlighted no taxonomic dominance across world oceans, according to the taxonomic composition associated to each cluster, suggesting a worldwide functional redundancy in the genomic potential supporting C4-enzymes (**Fig. S7**).

### 2.3. Weighted distribution of the genomic potential related to C4-photosynthesis

Here we present projections for each C4-enzyme and RUBISCO. First, we rescaled the cluster-level projections (i.e., model outputs; **Fig. S1D**) by their observed metagenomic read abundance (i.e., weighted distribution patterns, **Fig. S2**). Then, we aggregated these patterns at the enzyme-level according to their respective functional annotation. We therefore propagate the observed dominance of a given cluster (i.e., and associated taxa) to the aggregated enzyme-level patterns. The resulting enzyme-level projections are referred to as weighted patterns. For each enzyme, it represents the corresponding genomic potential (i.e., relative to the other considered enzymes), according to the environmental conditions at each geographical location.

We predicted contrasted weighted patterns between RUBISCO and across acid decarboxylation type (**Figure 3A**). Indeed, the weighted pattern of RUBISCO presented maximum potential in temperate areas (**Figure 3B**).

**Figure 3.**
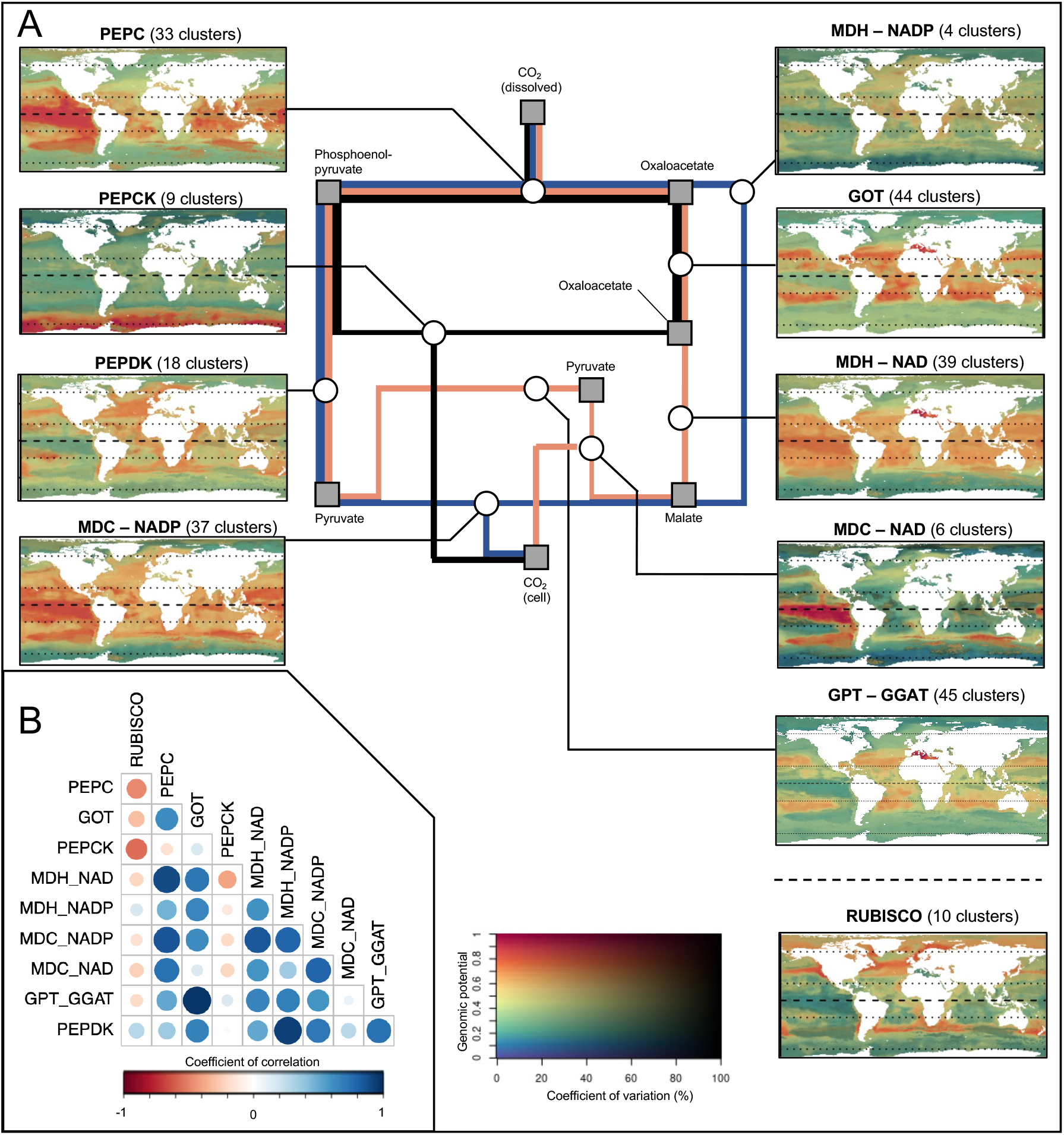
Weighted patterns corresponding to the relative genomic potential supporting C4-enzymes and RUBISCO, re-scaled by the corresponding observed relative metagenomic reads abundance. **(A)** Synthetic diagram of the metabolic pathway and corresponding projections. **(B)** Inter-projections Pearson’s spatial correlation index. The three mains currently described acid-decarboxylation types are represented in blue (Malate-NADP), red (Malate-NAD) and black (PEPCK), respectively. Involved metabolic components and enzymes are indicated on the diagram by squares and circles, respectively. The 2D color scale represents the weighted genomic potential for the target enzyme as the hue value (Y-axis) and the associated coefficient of variation as the saturation (i.e., uncertainty in % of the mean; X-axis). An orange to red hue corresponds to region where environmental conditions yield a high proportion (>0.6) of the target genes in the model. A low saturation level corresponds to an important variance among the underlying cluster-level projections.

We predicted low-to-moderate potential (< 0.3) and moderate (∼ 30 %) uncertainty in high latitudes for the weighted patterns of PEPC, MDCs, MDHs, and transferases (i.e., GOT and GPT – GGAT; **Figure 3A**). These patterns also presented moderate-to-high potential (between 0.5 and 1) in tropical areas, with some discrepancies. We show a Pearson’s correlation index above 0.5 between the above-mentioned enzymes, and above 0.7 for GOT and MDHs (**Figure 3B**). The latter presented an important potential in oligotrophic regions (e.g., Pacific gyres), suggesting functional redundancy in the genomic potential from Oxaloacetate to Malate (**Figure 3A**). In contrast, we predicted a high potential (> 0.7) in eutrophic Pacific waters for the weighted patterns of MDCs (Pearson’s correlation above 0.7; **Figure 3A**). Overall, we show high confidence in the areas associated to high genomic potential, with coefficient of variations lower than 30 % among all trained algorithms and 100-bootstrap projections. The above-mentioned weighted responses to environmental variables are similar to the ones highlighted in section *3*.*1*., characterized by higher potential in warm, low seasonality, and generally oligotrophic water bodies (**Fig. S4** and **S6**).

Conversely, we predicted moderate to high intensity values in oligotrophic tropical areas, but most importantly in the Southern Ocean (> 0.5; **Figure 3**) for the weighted pattern of PEPCK (i.e., a different acid decarboxylation type). The latter was preferentially distributed along water bodies characterized by (i) high seasonality of the Chlorophyll *a* concentration and the depth of the euphotic zone, (ii) high concentrations of oxygen (presenting the highest explanatory power in the model training; **Fig. S4**) and nutrients (e.g., phosphates and nitrates) and (iii) average temperatures below 8 °C (**Fig. S6**).

Finally, we highlighted that weighted patterns associated with high latitudes (e.g., correlated with the one of PEPCK) were composed at 28 % of Prymnesiophyceae and 50 % of Mamiellophyceae (Shannon index of 1.5), based on the taxonomic composition of each cluster. Mamiellophyceae also composed 40 % of the patterns with a clear temperate affinity (e.g., correlated with the one of RUBISCO; **Fig. S8**). In contrast, we highlight a larger diversity of taxonomic classes, with a Shannon index of 2.1, for patterns associated with equatorial latitudes.

## DISCUSSION

### 3.1. Genomic potential for C4-CCM in picoeukaryotes

By selecting clusters (i.e., PFCs) annotated by C4-enzymes or RUBISCO only, we considered a fraction of the available metagenomic information (i.e., ∼67 % of the clusters related to C4-enzymes or RUBISCO). In addition, genes related to other metabolic pathways may have responses to environmental variables different from genes related to C4-enzymes, potentially including bias in their corresponding PFC’s projection. Therefore, selecting a reduced set of clusters alleviates the risk of metabolic noise in the environmental responses, limited to the effect of C4-enzymes potentially involved in other pathways (e.g., GPT-GGAT transporter).

Our study focused on planktonic picoeukaryotes, the photosynthetic fraction of which is generally dominated by the Mamiellophyceae, Prasinophyceae, Prymnesiophyceae, Bacillariophyceae, and Dinophyceae lineages in the open ocean (16, 20). The potential for C4-photosynthesis has been suggested for several families, including Bacillariophyceae by combining C4-enzyme inhibition and photosynthetic efficiency monitoring (e.g., PEPDK 36, PEPC and PEPCK, 37). Evidence for genes encoding all C4-enzymes exist in *Micromonas* and *Ostreococcus*, Mamiellophyceae (38, 39). A plastid PEPC enzyme was recently discovered in *Emiliana huxleyi* (38), a Prymnesiophyceae abundant in temperate and polar regions (40). However, to our knowledge, no study provided univocal evidence for C4-CCM usage in situ. Stable isotope measurements would be necessary to fully understand C4-photosynthesis in picoeukaryotes, but they are difficult to apply at species-level in natural, uncultured, plankton communities (e.g., 8, 10). Alternatively, recent literature suggests the need for further studies on deep chlorophyll a maxima and various transporters (e.g., bicarbonate transporters), some of which are associated with or specific to C4 metabolism, to better understand C4-CCM in natural populations (5, 6).

Complementing these experimental approaches, we use a data-driven approach to shed more light on the environmental drivers of C4-genes in marine picoeukaryotes. However, MAGs integrate chloroplast and mitochondrial genes corresponding to C4-enzymes but do not distinguish their origin (28), nor provide information on the subcellular location of the corresponding enzymes (9, 41). Therefore, the patterns presented here must be interpreted as the potential for the (co-) presence of those pathways in the genome. They should be complemented by culture-based studies, locating enzymes within cells and/or performing carbon isotope discrimination to confirm C4-CCM presence, expression, and its co-existence with C3-photosynthesis in picoeukaryote lineages (8). The present study could be used to locate regions where such mechanisms are most likely to occur.

### 3.2. Environment-driven genomic potential

The modeled distribution patterns revealed that the genomic potential for C4-photosynthesis is more associated with tropical oligotrophic and annually stratified waters. Conversely, the proportion of reads related to RUBISCO (i.e., considered as a representative of all photosynthetic pathways, due to its central role in C3, C4 and CAM photosynthesis) is higher in temperate regions (**Figure 2A**). The fact that terrestrial C4-plants (4) and the genomic potential for C4-CCM in picoeukaryotes display similar latitudinal distribution, around the tropics, does not imply that the environmental drivers of those distributions are the same. In terrestrial plants, C4-CCMs are considered as an adaptation to drought and are, for example, also associated with a specific leaf structure that reduces their water consumption (4). Drought is of course not an evolutionary driver for marine picoeukaryotes. Alternatively, they present an important surface-to-cytoplasm ratio (i.e., small cells or presence of a vacuole, 42, 43) leading to a high nutrient absorption yield, which is adapted to oligotrophic waters, common in the tropical ocean.

In addition to environmental conditions, the biogeography of the genomic potential supporting C4-CCM may also relate to irradiance levels, largely controlling ATP generation, necessary to the decarboxylation reaction (42). Indeed, C4-CCM requires additional ATP generation to increase the RUBISCO efficiency in comparison to classical C3-photosynthesis, without impacting the energy available for the latter (42, 44). In contrast, an excess of ATP may lead to photoinhibition, thus lower carbon fixation efficiency (36, 45). Therefore, it has been suggested that C4-photosynthesis is particularly adapted to dissipate excess energy in the cell in high irradiance areas such as tropical oceans (5, 36). Our weighted patterns highlighted differences between PEPCK and MDCs (**Figure 3**). The latter require 2 extra ATP compared to the C3 carbon fixation to complete the pathway. In a logical way, the PEPCK acid decarboxylation type, which only requires 1 extra ATP and thus is supposed to be more efficient in low irradiance environments (44), showed here the highest genomic potential in polar or sub-polar regions.

### 3.3. Functional and ecological implications

We highlighted functional redundancy among C4-genes in oligotrophic tropical waters (**Fig. S7**). This contrasts with high latitudes, where only a few taxa dominate (**Fig. S8**) (17, 46). More interestingly, we highlighted a biogeographical differentiation between the weighted pattern of RUBISCO – i.e., the baseline photosynthetic enzyme – and those of C4-enzymes. Since 30 million years ago, atmospheric CO_2_ concentration has drastically reduced from c.a. 1000 ppm to less than 200 ppm 20.000 years ago, resulting in lower dissolved carbon in the oceans (4). This led to a selective pressure towards efficient photosynthetic metabolism, like C4-photosynthesis (7) or, in a lesser extent, RUBISCO of higher carboxylation affinity (e.g., type II in Dinoflagellates, 9). While the evolution of C4-CCM in marine organisms is not yet fully understood, 48 independent evolutions of C4-CCM were identified in the genome of terrestrial plants (e.g., grasses, caryophyllales, 4), suggesting a higher genomic potential for C4-photosynthesis in taxonomically diverse areas (7). The above-mentioned functional redundancy in the genomic potential for C4-CCM in taxonomically rich tropical waters may relate to a co-evolution between taxonomic diversification and its associated functions (i.e., neutral theory). However, the functional diversity among C4 acid-decarboxylation types may also reflect – or be amplified by – a selection process, as it may present a selective advantage. Moreover, the respective dominance of Mamiellophyceae in temperate latitudes (i.e., correlated with the patterns associated to RUBISCO) and Prymnesiophyceae in polar latitudes, are concordant with the literature (40, 47), thus validating the environmental predictors controlling their biogeography. We identified key environmental predictors shaping the biogeography and (co-)dominance patterns of the genomic potential supporting C4-enzymes and RUBISCO. Such results open new perspectives of exploring the relationship between functional and taxonomic diversity in the oceans, complementing already diverse approaches and data types, and better understand the environmental drivers of key biogeochemical cycles in the current and future climatic context.

## MATERIAL AND METHODS

### 4.1. Data

#### 4.1.1. Genomic data

We studied the biogeography of the genomic potential related to C4-CCM through the prism of Metagenomic Assembled Genome (MAG, 28) retrieved from the *Tara Oceans* expedition (2009-2013). Briefly, 280 billion reads from 798 metagenomes, corresponding to the surface and deep chlorophyll maximum layer of 210 stations from the Pacific, Atlantic, Indian, Southern and Arctic Oceans, as well as the Mediterranean and Red Seas (**Figure 1**), encompassing eukaryote-enriched plankton size fractions ranging from 0.8 μm to 2 mm, were used as inputs for 11 metagenomic co-assemblies (6–38 billion reads per co-assembly) using geographically bounded samples. We thus created a culture-independent, non-redundant (average nucleotide identity <98%) genomic database for eukaryotic plankton in the sunlit ocean consisting of 683 MAGs and 30 single-cell genomes (SAGs), all containing more than 10 million nucleotides for a total size of 25.2 Gbp and encoding for 10,207,450 genes. Then, a sequence similarity network was built out using the 683 manually curated MAGs following a similar methodology to the one developed in Faure et al. (32). A pairwise comparison was computed between each protein sequence. The resulting alignment was then filtered, removing self-hits and pairs showing less than 80% of sequence identity and coverage. Resulting Protein Functional Clusters (PFCs, as in 32) were built, hereafter referred to as clusters. The functional annotation performed with eggNOG mapper v2.1.5 was added on the sequences, and the functional homogeneity was checked in each cluster (48, 49). The surface and metagenomic samples correspond to 130 stations.

#### 4.1.2. Environmental data

For each of the 130 selected *Tara Oceans* metagenomic surface samples, we retrieved a set of monthly, global scale, environmental climatologies encompassing the 2005 to 2017 period, at a spatial resolution of 1° x 1° (**Table S2**). The latter corresponds to the available climatology encompassing the sampling period (2009-2013), where we considered temporal environmental variations negligible in comparison to spatial environmental gradients. They correspond to a restricted set of factors characterizing the water body (e.g., oligotrophic, eutrophic) and related to C4-photosynthesis, for which we calculated the yearly average and yearly standard deviation (*i*.*e*., proxy of seasonal variations).

### 4.2. Data selection and pre-processing

#### 4.2.1. Protein functional cluster selection

We first selected a reduced set of clusters, within the 0.8 to 5 μm size fraction and surface samples, for which 100% of the KEGG Orthology (KO, 50) annotated protein members were related to C4-enzymes or RUBISCO (**Fig. S1, Table S1**). To avoid model over-parameterization and because rare clusters were assumed as not influencing the large-scale patterns investigated in this study, we only considered clusters that were present in a minimum of 10 *Tara Oceans* stations.

The corresponding dataset contained 240 clusters distributed across 130 *Tara Oceans* stations. The 240 clusters, functionally annotated with C4-enzymes and RUBISCO, were associated with 234 MAGs. The latter presented an average completeness estimate of 57% (**Table S3**). In comparison, the average completeness estimate across all MAGs from Delmont et al. (28) yield at 37 %. As a supplementary quality check, we estimated a minimum horizontal coverage (i.e., number of bases of a MAG covered with a certain depth) of 68 % for each of the 234 MAGs (**Table S3**). Finally, we show that our MAGs are associated with an average BUSCO completeness (i.e., the percentage of mapped BUSCO genes in each MAG) of 55.7% (**Table S3**). We therefore consider these MAGs of sufficient quality for identifying C4-genes across our samples.

To reduce the number of response variables (clusters; PFCs) to a reasonable amount for multivariate modelling, with respect to the limited number of stations, we performed an Escoufier dimensional reduction (51). The latter iteratively selects the clusters whose pattern across stations minimize the residual variance of the dataset. Here, we selected 50 clusters that represent over 95% of the 240 clusters variance to be included in the multivariate algorithm.

#### 4.2.2. Metagenomic data pre-processing

Genes abundances among samples were determined by mapping raw metagenomic reads against the gene database (28). Briefly, reads were mapped using the bwa tool, and only random best matches with at least 95% of sequence identity over at least 80% of the read length were retained as positive. To alleviate the effect of gene length and sequencing effort variability between samples on the number of reads, we normalized the metagenomic reads by the length of the corresponding gene coding part and the total number of reads per station (i.e., including reads of all non-considered clusters), respectively. Because the total genomic material present at each sampling station is unknown (i.e., non-exhaustive sampling and sequencing effort), the absolute number of reads is not comparable among stations. To compare the abundance between selected clusters at different sampling stations, we transformed the dataset to relative abundance (**Supplementary information text** and **Fig. S1**).

### 4.3. Multivariate Boosted Regression Tree

#### 4.3.1. General principle

Recently, growing interest for interactions between response variables led to the development of multivariate machine learning algorithms, such as Multivariate Boosted Tree Regressors (MBTR, 35). The latter is also particularly adapted to small sample size as the interactions between response variables is considered as supplementary information to calibrate the model. Here, MBTR is used to model the relationship between climatologies and metagenomic relative abundance (i.e., summed at 1 for each station; **Supplementary information text** and **Fig. S1**). To best reproduce the response of metagenomic reads (i.e., response variable) to the corresponding environmental variables (i.e., explanatory variable), the model sequentially fits decision trees (i.e., boosting rounds) using gradient descent to minimize a specific loss function (see **Supplementary information text** for hyperparameter and loss choice). At each boosting round, the algorithm fits a decision tree on the residuals of the previous boosting round and computes a tree loss (i.e., a measure of deviation between observed and predicted response variable values). Decision trees are constructed using the hessian of the loss function (i.e., second order tensor of its partial derivatives) to minimize the loss gradient. Therefore, the information learned by the *n*^*th*^ tree is passed to the *n+1*^*th*^ tree at a user-defined learning rate (**Supplementary information text** and **Fig. S1**). The ensemble of sequentially fitted decision trees are considered in the model until the minimum loss is reached. Finally, one important feature of MBTR is the conservation of the initial correlation structure between the response variables (see methods in 35). The latter is tested by computing a Pearson correlation matrix between response variables before and after model fitting, whose conservation is tested by a Mantel matrix comparison test (**Supplementary Information text**).

#### 4.3.2. Model training and evaluation

To avoid over-fitting, the explanatory and response datasets were split between training set and test set using a *n*-fold cross-validation procedure. For each model, *n* algorithms were trained on different *n-1* folds, while the remaining fold was used for testing only (i.e., computing the loss at each boosting round). To minimize the effect of spatial and temporal autocorrelation in our data (i.e., leading to over-optimistic model evaluation, 52), the *n*-folds were defined according to the *Tara Oceans* station number. Because the cruise followed a continuous trajectory in time and along the sampled stations, the resulting folds are spatially and temporally distant (i.e;, spatial and temporal block splitting, as recommended in 52). The resulting *n*-algorithms predictions were aggregated in an average response and its corresponding coefficient of variation (CV). The ability of the final model to reproduce the observed clusters relative abundance across environmental conditions has been measured by the R^2^ criteria and the root mean square error (RMSE, between 0 and 1 according to the distribution pattern scale).

#### 4.3.3. Spatial projections

To better estimate projection uncertainty, our spatial projections were constructed using a bootstrap procedure. For each 100-bootstrap round, we first re-sampled the original dataset (i.e., train and test response dataset and corresponding explanatory variable values) with replacement. Then, we re-fitted an MBTR algorithm on the re-sampled data by using the hyperparameters corresponding to the validated model, including the number of boosting rounds corresponding to the minimum loss across all *n*-algorithms. Finally, the re-fitted MBTR algorithm was used to predict the relative abundance of clusters worldwide, using the corresponding climatologies values at each geographical cell.

### 4.4. From model projections to final outputs

We only modelled the 50 clusters representing 95% of the dataset variability. Therefore, we indirectly reconstructed the projections of the 190 others by identifying their most representative Escoufier-selected cluster. To this extent, we performed a correspondence analysis based on the observed relative abundance of all clusters. By using the dimensions of the correspondence analysis space corresponding to a minimum of 80% variance explained, we calculated the Euclidean distance between each non-selected cluster, and its nearest neighbor selected by the Escoufier criteria. Because the 50 Escoufier selected clusters represented over 95% of the dataset variability, we considered that a cluster and its nearest neighbor in the correspondence analysis space share the same relative abundance pattern. In addition, we calculated the scale of each non-selected cluster with respect to their nearest Escoufier-selected neighbors using the sum of their observed relative abundance across all stations (**Fig. S2**). We then reconstructed the spatial projections of the 190 clusters not considered in MBTR according to their projected nearest Escoufier-selected neighbor. The resulting 240 cluster-level projections of the genomic potential were then aggregated at the enzyme level according to their functional annotation (see Result section, **Fig. S2**).

## Acknowledgments

The co-authors wish to thank public taxpayers who fund their salaries. A.S. and P.D.’s salaries were financed by the Blue-Cloud European project (Grant Agreement n.862409). The authors want to thank all people involved in the Tara Oceans project for making data publicly available. Finally, the authors wish to thank the editor and the three anonymous reviewers for their helpful comments on this manuscript.

## Author Contributions

S.D.A., L.G. and J.O.I. conceived the study. L.G. and J.O.I. supervised the study. P.D., L.B. and E.P. processed the metagenomic data and provided expert advice on their use and interpretation. A.S. wrote the first draft, the modelling pipeline and performed the analysis. All authors substantially contributed to the successive versions of this manuscript.

## Data availability

Instructions on how to build the Sequence Similarity Network and associated Protein Functional Clusters are available at: https://data.d4science.net/BN9t. Instructions and credentials on how to access the genomic database used in this study (PostgreSQL) are available in the technical documentation at https://data.d4science.net/qa7Z.

The ensemble of enzyme-level projections are available upon registration at [https://data.d4science.net/Zraq]. Additional data that support the findings of this study are available from the corresponding author upon reasonable request.

## Code availability

All R and Python codes, the corresponding pipeline, libraries, and associated technical documentation are available in the Blue-Cloud catalogue at: https://data.d4science.net/qa7Z

