## Supplementary material for "Picoeukaryotic photosynthetic potential is functionally redundant but taxonomically structured at global scale": Fig. S

*Corresponding author

**These authors contributed equally

### SUPPORTING INFORMATION TEXT

#### 1. Supplementary methods

This supplementary material contains additional details on the framework developed for this study, that is synthesized in **Fig. S1 and S2**.

##### 1.1. Metagenomic data construction

Our study focused on 683 eukaryotic Metagenome-Assembled-Genomes (MAGs). The whole bioinformatic workflow designed to build these MAGs can be found in the preprint as well as the supplemental information on the genoscope website:

(https://www.biorxiv.org/content/10.1101/2020.10.15.341214v2)
(https://www.genoscope.cns.fr/tara/)

These were the first eukaryotic MAGs from a comprehensive genome-resolved metagenomic survey using 798 metagenomes derived from the *Tara Oceans* expeditions. They correspond to the surface and deep chlorophyll maximum layer of 210 stations from the Pacific, Atlantic, Indian, and Southern Oceans, as well as the Mediterranean and Red Seas, encompassing eight eukaryote-enriched plankton size fractions ranging from 0.8 µm to 2 mm). Following read recruitment of the metagenomes and metatranscriptomes the MAGs were organized into four major size classes of 0.8 to 5 µm, 3 to 20 µm, 20 to 180 µm and 180 to 2000 µm. Original metagenomes are available under the European Bioinformatics Institute (EBI) repository with project ID PRJEB402 (see Table S1 in Delmont et al., 2022 for detailed reports of accession numbers and additional information for each metagenome). To date, the work achieved by Delmont et al. (1) constitutes the only database of manually refined MAGs constructed using the Tara Oceans project data.

To estimate the abundance and expression of each contig in each sample, cleaned reads (from metagenomes and metatranscriptomes) were mapped against the eukaryotic MAGs using the bwa tool (version 0.7.4). The following parameters were used: bwa aln -l 30 -O 11 -R 1; bwa sampe -a 20000 -n 1 –N; samtools; rmdup. Low complexity reads were removed. Reads covering at least 80% of read length with at least 95% of identity were retained for further analysis. In the case of several possible best matches, a random one was picked.
A first sequence similarity network was built out of 683 manually curated MAGs from 10,207,435 eukaryotic proteins. This file was used for the creation of a diamond database and a protein blast of the protein sequences against the database to compute the percentage of similarity between every pair of proteins detected in the MAGs. Here we used a maximum e-value of 1e-3, and the sensitive option adapted to long reads (−e 1e−3 −p 30–sensitive).
The alignment was then filtered removing all self-hits. Several thresholds for percentage of identity and coverage were tested (75%, 80%, 85% and 90%). A Sequence Similarity Network (SSN) was built with the diamond output using 80% identity and 80% coverage threshold to minimize the number of singletons while maximizing the functional homogeneity between linked proteins (reproducible analysis and statistical exploration provided in Meng et al. (2), Rizzolo et al. (3), Faure et al. (4)).
An SSN is made of singletons (vertices or sequences without any homology with other sequences) and connected components (CC; i.e., subgraphs composed of at least two vertices disconnected from the rest of the network). In our case, a CC corresponds to a group of at least two protein sequences that are linked together (directly or via neighbors), and that have no link with other groups of sequences in the SSN. We assume that the proteins contained in a CC potentially share a similar molecular function (2, 3, 5, 6). Our SSN was composed of 1,287,251 CCs, including 4,749,034 proteins. These proteins were functionally annotated using eggNOG mapper v2.1.5.

1.2. Data selection and pre-processing

The first step of our modelling pipeline (**Fig. S1A**) consisted in selecting a set of protein functional clusters (PFCs; hereafter referred to as clusters) markers of the C4-carbon concentration pathway enzymes as well as the baseline photosynthetic enzyme (i.e., the RUBISCO). The corresponding selected enzymes are described in **Table S1.**

We then extracted the metagenomic reads relative abundance per *Tara Oceans* station and clusters from the SSN presented in the previous section. Note that starting from station n°66 (i.e., Cape Town), a new size class of 0.8 – 2000 µm has been implemented in the *Tara Oceans* cruise, while the initial 0.8 – 5 µm was not sampled from stations 155 onwards (i.e., Arctic stations). Given that smaller organisms are much more abundance than large ones the organisms sampled with the 0.8 – 2000 µm filter are essentially picoeukaryotes. And indeed, the composition of metagenomic reads for the clusters of interest at common stations (i.e., 66 to 155) showed significant correlation (Pearson correlation: 0.89; p-value: 0.01) between the two filters, supporting the inclusion of Arctic data issued from the 0.8 – 2000 µm filter.

As mentioned in *section 4.2.1*. of the manuscript, the effect of the different selection criterion such as the exclusivity and the minimum number of stations coverage are shown in **Fig. S2**. The main loss of signal is observed for K00028 and K00029 enzymes, corresponding to Malate dehydrogenase (decarboxylating; NAD and NADP) as well as the Prymnesiophyceae taxonomic class. All selection criteria resulted in the same distribution as the baseline data (i.e., all PFCs related to either RUBISCO or C4 enzymes), except for the KO composition, due to the above-mentioned loss of signal.

1.3. Model training, evaluation, and projections

The second step of our modelling pipeline (**Fig. S1B**) corresponded to the model training on 50 clusters, whose patterns were best representing the dataset variability.

First, we fitted one Multivariate Boosted Tree Regressor (MBTR) algorithm per training set and hyperparameter combination, computing the loss at each boosting round. The MBTR algorithms were fitted under a mean square error loss function, a learning rate of $5.10^{-3}$ and a 10-fold cross validation procedure. Additionally, we set the minimum number of observations in terminal leaves to 30 and the number of quantiles considered to find the best split to 10. The model loss is adapted to discriminate between different sets of hyperparameters. Thus, we only considered the set of hyperparameters that resulted in the minimum loss across the corresponding 10 trained MBTR algorithms. However, the loss is not providing information on the actual performance of the model in reproducing the observed data. Therefore, for each of the 10 trained MBTR algorithms, we predicted the relative abundance of the metagenomic reads on the environmental values corresponding to each of the 10 test sets. The 10 corresponding predictions were compared against the truth (i.e., observed values) by means of the R-squared (R2) and Root Mean Square Error (RMSE, here between 0 and 1) to assess model performance on data not seen by the MBTR models during the training process. The corresponding model evaluation estimated a R2 of 0.33 and RMSE of 0.05. We also tested the conservation of the correlation structure between the response variable during the model training. In our case, the correlation structure is conserved with a mantel correlation value of 0.748 and a p-value of 0.01. Finally, for the spatial projections, we performed a total of 100 bootstrap rounds and computed the average and CV between all bootstrap-projections (**Fig. S1**).

### 2. Variable importance and partial dependence plots

This supplementary material presents the variable importance in the model training (**Fig. S3**). The latter is calculated as the number of times an environmental variable was selected for a tree split, scaled by the corresponding loss gain.

Following model evaluation and variable importance, we calculated the response of the genomic potential of each enzyme, to each environmental variable, in the form of partial dependence plots. The latter are represented in **Fig. S5** and **S6**, respectively corresponding to the standardized and weighted patterns.

#### 3. Estimating the taxonomic effect on the distributional patterns of genomic potential

To estimate the taxonomic composition associated with each pattern (i.e., standardized or weighted), we constructed the distribution pattern of each MAG. We then performed a hierarchical clustering to construct projections of MAGs of similar patterns that were then correlated to the enzyme's distributional patterns. Finally, according to each MAG annotation, we computed the taxonomic composition corresponding to each cluster of MAGs patterns. The results are presented in **Fig. S6** and **S7**, respectively for the standardized and realized patterns.

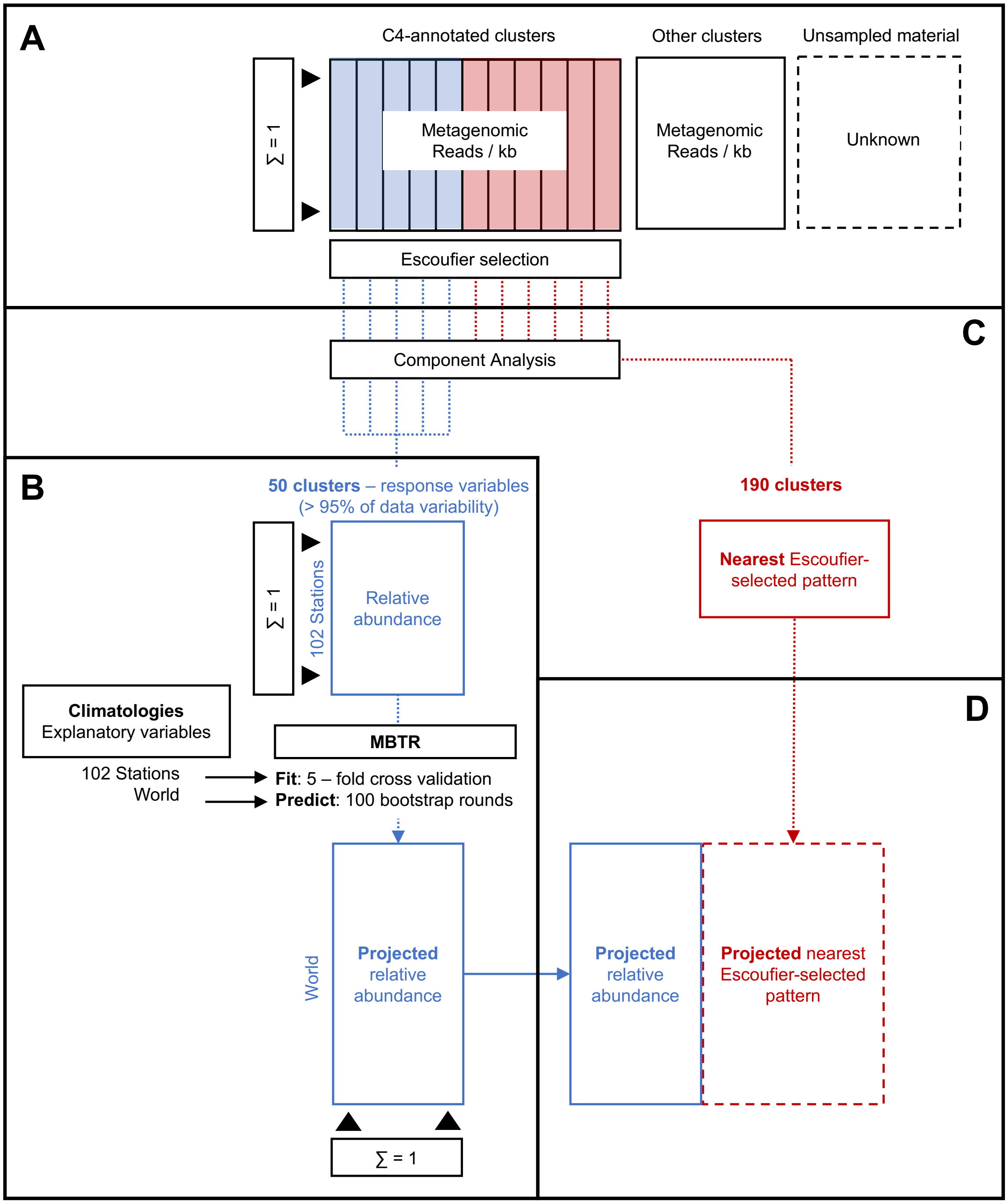


**Fig. S1.** Synthetic diagram presenting the successive modelling steps, including (A) the genomic data selection and pre-processing, (B) the Multivariate Boosted Tree Regressor (MBTR) training and projections, (C) the consideration of protein functional clusters not selected in MBTR and (D) the resulting projections at the cluster-level.

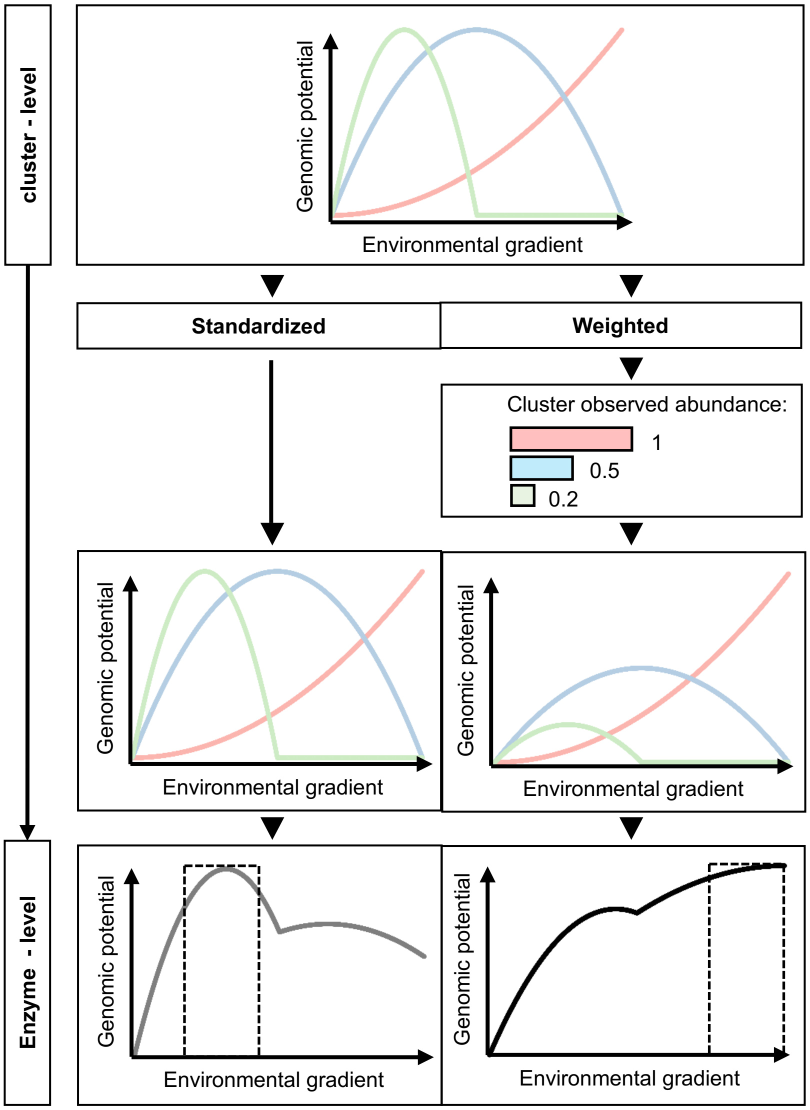


**Fig. S2:** Synthetic diagram describing the pattern aggregation methods from the cluster-level to the enzyme-level. The left panels represent the construction of standardized patterns, where all cluster-level patterns are aggregated with equal weight into the enzyme-level pattern. The right panels represent the construction of weighted patterns, where each cluster-level pattern is weighted according to its observed abundance when aggregated into the enzyme-level pattern. The dashed box in the bottom panels represents the highest genomic potential corresponding to each aggregation method, with respect to the environmental gradient.

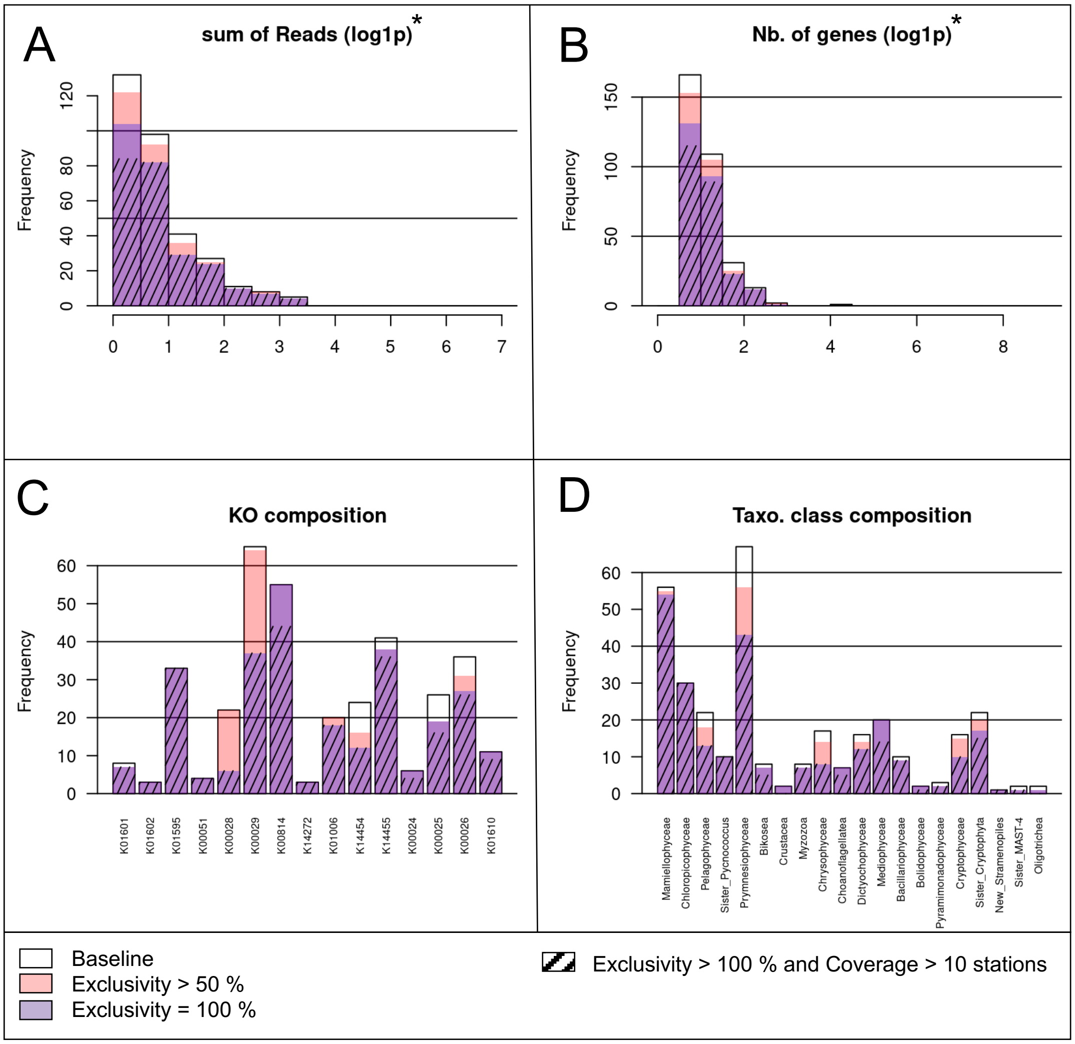


* all filters have significantly the same distribution as "Baseline" with 95% significance

**Fig. S3.** Effect of various protein functional cluster selection criteria on their composition.

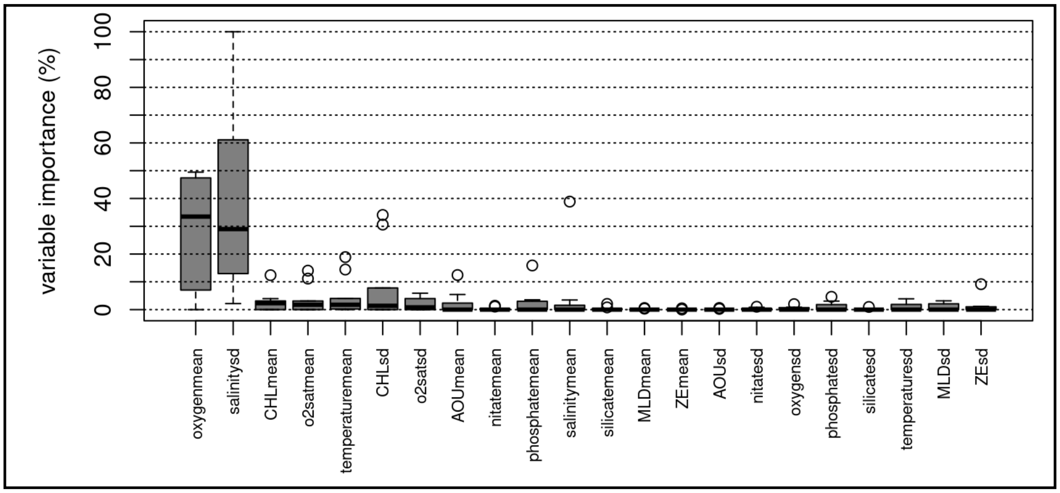


**Fig. S4.** Environmental variable importance in model training, across cross-validation runs. The midline of the boxplots corresponds to the median, with the upper and lower box limits corresponding to the first and third quartile. Whiskers extend to 1.5 times the interquartile range and white points correspond to outliers.

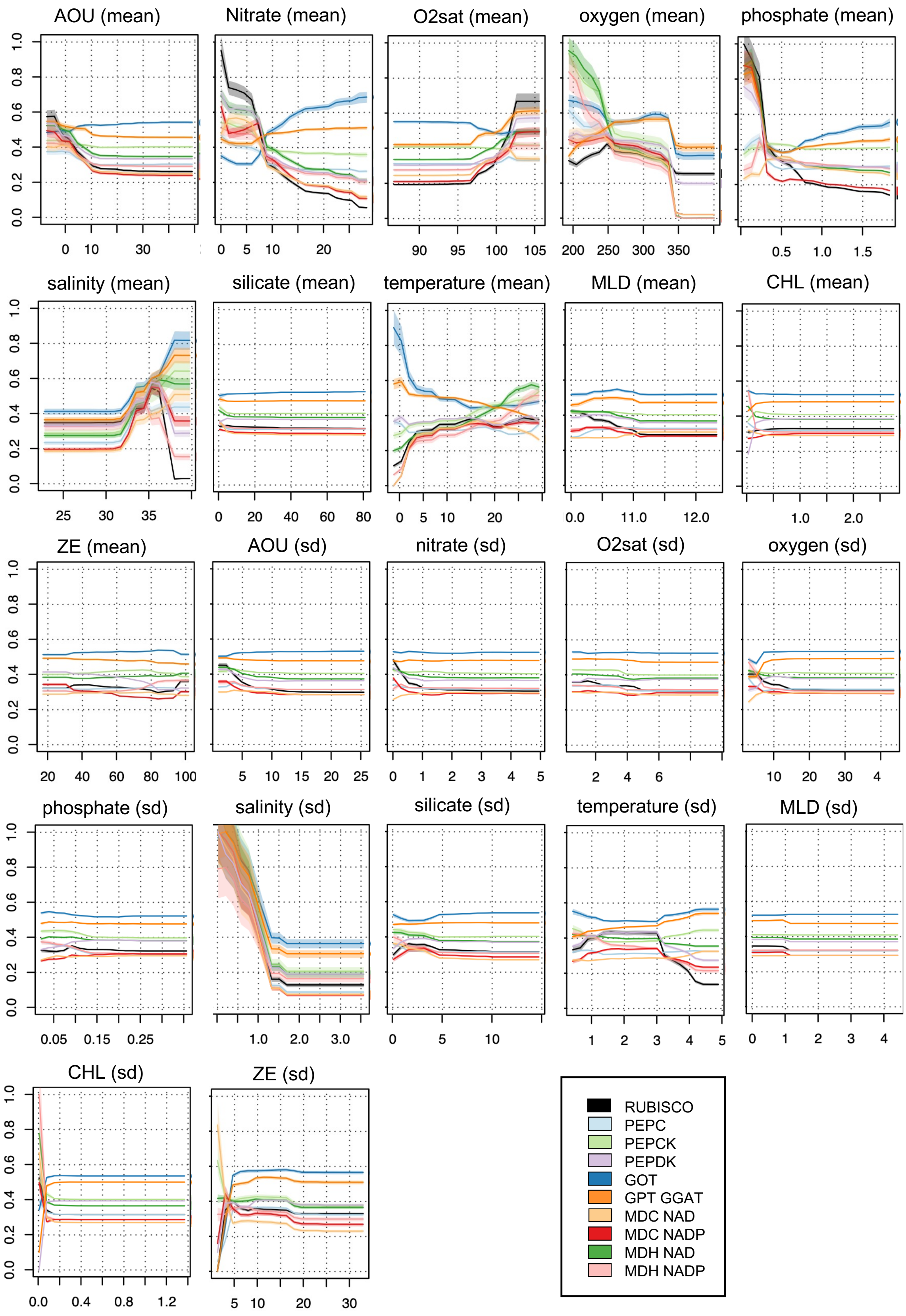
AOU (mean)
Nitrate (mean)
O2sat (mean)
oxygen (mean)
phosphate (mean)
salinity (mean)
silicate (mean)
temperature (mean)
MLD (mean)
CHL (mean)
ZE (mean)
AOU (sd)
nitrate (sd)
O2sat (sd)
oxygen (sd)
phosphate (sd)
salinity (sd)
silicate (sd)
temperature (sd)
MLD (sd)
CHL (sd)
ZE (sd)
0.0
0.2
0.4
0.6
0.8
1.0
0 10 30
0 5 10 20
90 95 100 105
200 250 300 350
0.5 1.0 1.5
25 30 35
0 20 40 60 80
0 5 10 20
10.0 11.0 12.0
1.0 2.0
20 40 60 80 100
5 10 15 20 25
0 1 2 3 4 5
2 4 6
10 20 30 4
0.05 0.15 0.25
1.0 2.0 3.0
0 5 10
1 2 3 4 5
0 1 2 3 4
0.0 0.4 0.8 1.2
5 10 20 30
RUBISCO
PEPC
PEPCK
PEPDK
GOT
GPT GGAT
MDC NAD
MDC NADP
MDH NAD
MDH NADP

**Fig. S5.** Partial dependence plots corresponding to the standardized pattern. The Y axis corresponds to the genomic potential while the X axis represents the environmental parameter possible values across world oceans. The full line and shading respectively correspond to the average response and standard deviation between bootstrap runs.

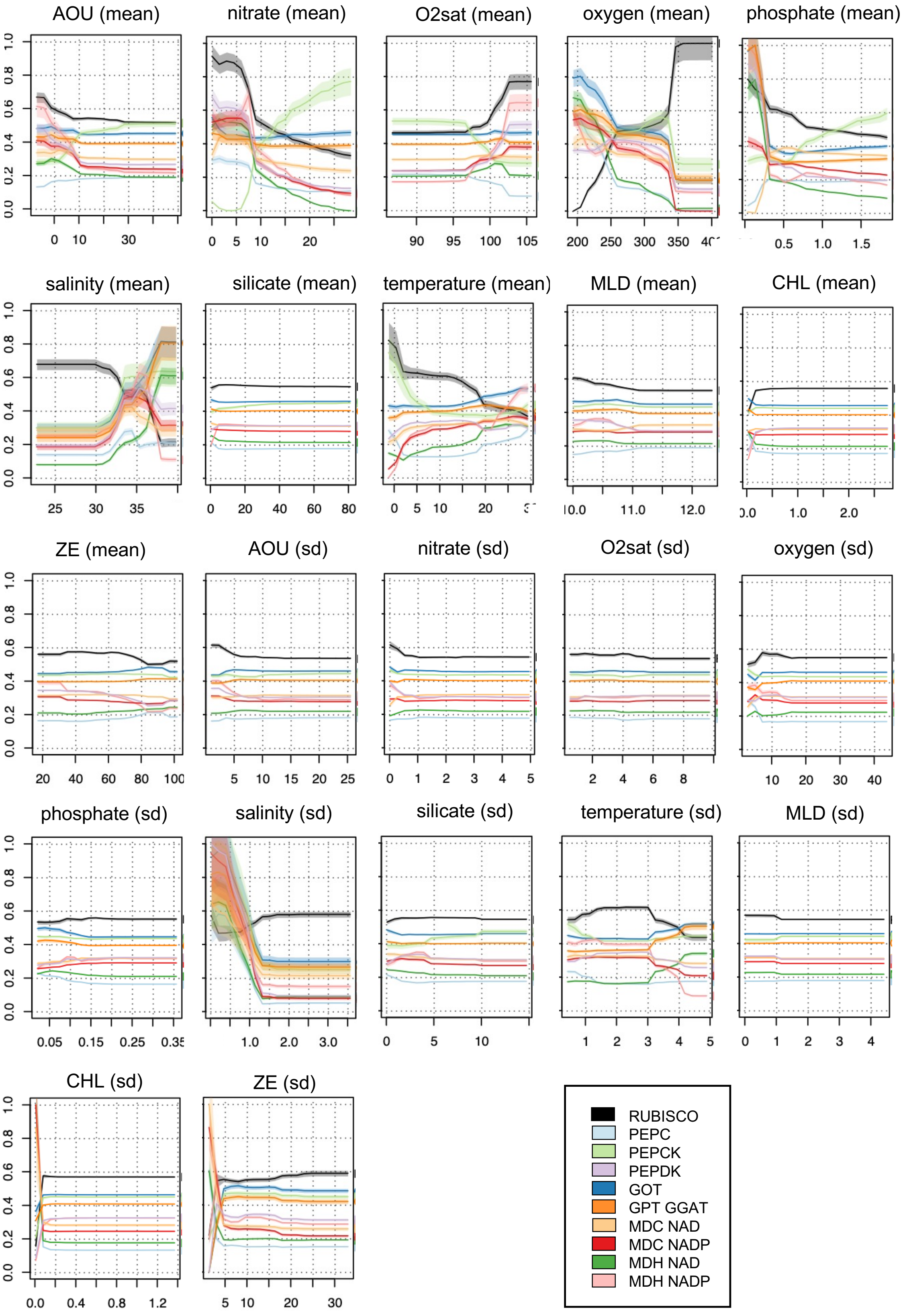
AOU (mean)
nitrate (mean)
O2sat (mean)
oxygen (mean)
phosphate (mean)
salinity (mean)
silicate (mean)
temperature (mean)
MLD (mean)
CHL (mean)
ZE (mean)
AOU (sd)
nitrate (sd)
O2sat (sd)
oxygen (sd)
phosphate (sd)
salinity (sd)
silicate (sd)
temperature (sd)
MLD (sd)
CHL (sd)
ZE (sd)
RUBISCO
PEPC
PEPCK
PEPDK
GOT
GPT GGAT
MDC NAD
MDC NADP
MDH NAD
MDH NADP

**Fig. S6.** Partial dependence plots corresponding to the weighted pattern (i.e., re-scaled by the corresponding observed relative metagenomic reads abundance). The Y axis corresponds to the genomic potential while the X axis represents the environmental parameter possible values across world oceans. The full line and shading respectively correspond to the average response and standard deviation between bootstrap runs.

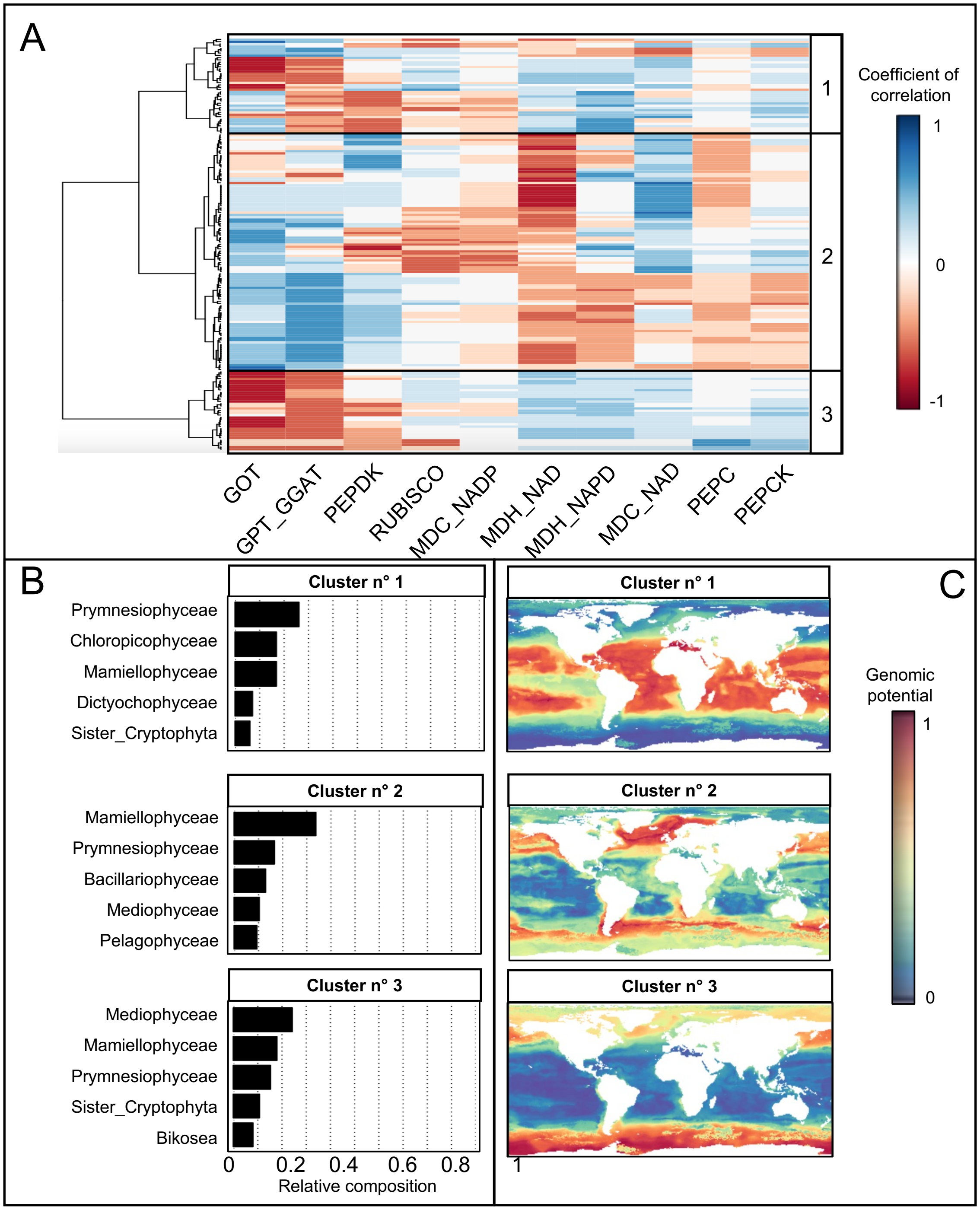


**Fig. S7.** Estimated MAG-based taxonomic effect on standardized patterns with (**A**) the correlation between MAG distribution pattern clusters and projected genomic potential related to a given enzyme, (**B**) the taxonomic composition of each MAG cluster and (**C**) their corresponding projections.

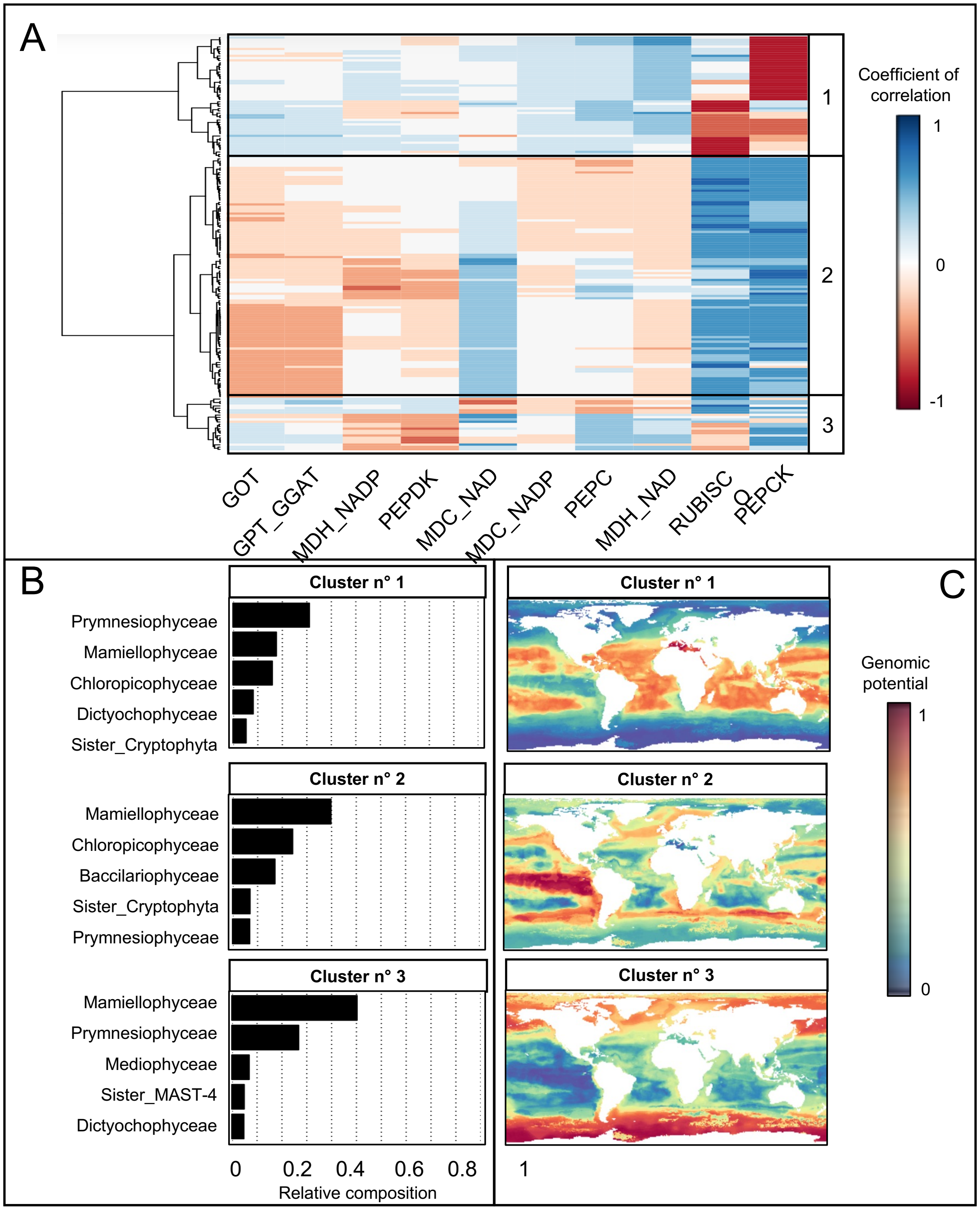


**Fig. S8.** Estimated MAG-based taxonomic effect on weighted patterns (i.e., re-scaled by the corresponding observed relative metagenomic reads abundance) with (**A**) the correlation between MAG distribution pattern clusters and projected genomic potential related to a given enzyme, (**B**) the taxonomic composition of each MAG cluster and (**C**) their corresponding projections.

**Table S1.** C4 carbon concentration-related enzymes and corresponding KEGG Orthology (KO) and enzyme (EC) reference annotations.

| **Enzyme** | **Description** | **KO** | **EC** |
|---|---|---|---|
| PEPC | Phosphoenolpyruvate carboxylase | K01595 | 4.1.1.31 |
| GOT | Aspartate aminotransferase, cytoplasmic | K14454 | 2.6.1.1 |
| | Aspartate aminotransferase, mitochondrial | K14455 | |
| PEPCK | Phosphoenolpyruvate carboxykinase (ATP) | K01610 | 4.1.1.49 |
| MDH – NADP | Malate dehydrogenase (NADP) | K00051 | 1.1.1.82 |
| MDH – NAD | Malate dehydrogenase (NAD) | K00024; K00025; K00026 | 1.1.1.37 |
| MDC – NADP | Malate dehydrogenase (decarboxylating; NADP) | K00029 | 1.1.1.40 |
| MDC – NAD | Malate dehydrogenase (decarboxylating; NAD) | K00028 | 1.1.1.39 |
| GPT - GGAT | Alanine transaminase | K00814 | 2.6.1.2 |
| | Glutamate – glyoxylate aminotransferase | K14272 | 2.6.1.4; 2.6.1.44 |
| PEPDK | Pyruvate, orthophosphate dikinase | K01006 | 2.7.9.1 |
| RUBISCO | Ribulose-1,5-biphosphate carboxylase oxygenase | K01601 K01602 | 4.1.1.39 |

**Table S2.** Environmental climatologies considered in the model.

| Name | Description | Reference |
|---|---|---|
| temperature | Sea surface temperature (°C) | Boyer et al. (7) |
| salinity | Sea surface salinity (unitless) | |
| oxygen | Dissolved Oxygen (μmol/kg) | |
| o2sat | Percent Oxygen Saturation (%) | |
| AOU | Apparent Oxygen Utilisation (μmol/kg) | |
| silicate | Silicate (μmol/kg) | |
| phosphate | Phosphate (μmol/kg) | |
| nitrate | Nitrate (μmol/kg) | |
| ZE | Depth of the Euphotic Zone (m) | Morel & Maritorena (8) |
| MLD | Mixed Layer Depth (m) | de Boyer Montégut et al. (9) |
